# The Time Varying Networks of the Interoceptive Attention and Rest

**DOI:** 10.1101/840645

**Authors:** Ana Y. Martínez, Athena Demertzi, Clemens C.C. Bauer, Zeus Gracia-Tabuenca, Sarael Alcauter, Fernando A. Barrios

## Abstract

Focused attention to spontaneous sensations is a phenomenon that demands interoceptive abilities and a dynamic character of attentive processes. The lack of its control has been linked to neuropsychiatric disorders, such as illness-anxiety disorder. Regulatory strategies, like focused attention meditation, may enhance the ability to control attention particularly to body sensations, which can be reflected on functional neuroanatomy. Adopting a systems-level approach, we aimed at estimating the recurring fMRI functional connectivity (FC) patterns between regions of the dorsal attention network, default mode network, and frontoparietal network during 20 minutes of an attentional task to spontaneous sensations (Task), and at rest, before (Pre-task rs) and after the task (Post-task rs), in fifteen experienced meditators. Dynamic functional connectivity analysis was performed using sliding windows and k-means clustering on the grouped data finding five FC patterns. In both rest conditions the subjects remain longer in a low connectivity state, in contrast, during the task a higher proportion of time spent in complex organization states was preferred. Moreover, an impact over the post-task rs FC was observed as an effect of the preceding interoceptive task performance, with this remaining effect probably taking an active role in the learning process linked to cognitive training.

## Introduction

In our everyday life, numerous stimuli surround us, requiring their selection and processing through attention (Posner, 2012). Once a stimulus is attended, it will be more likely for it to influence the brain systems and to guide our behavior (Dehaene and Changeux, 2005; Webb and Graziano, 2015). Attention also exerts an important modulation of the body representation in the somantosensory cortex. It is associated with the facilitation of the conscious perception of the external stimuli and also can increase or decrease the perception of spontaneous sensations, occurring without any external stimulus, as a response of the focused attention on the body (Boly et al., 2007; Ferentzi et al., 2018). Spontaneous sensations are closely related to interoception, the sensing of the body’s physiological condition, essential for body awareness (Michael et al., 2015; Michael and Naveteur, 2011).

As a property of multiple cognitive processes (Bartolomeo and Chokron, 2000; Golomb and Turk-Browne, 2010), the lack of regulation in the control of attention is linked to different psychiatric disorders (Donald et al., 2014; White and Shah, 2006), like panic disorder, somatization and illness anxiety disorders which are distinguished by an excessive attention and increased concern in body sensations (Stern et al., 2017; Stins et al., 2015).

Regulatory strategies, like focused attention meditation (FAM) practices, considered a form of cognitive training (Tang et al., 2015), may enhance the ability to control the attention particularly to body sensations, producing an improvement in the attentional skills. FAM requires to focus the attention to an object and bring the attention back to the object when it is lost (Manna et al., 2010), resulting in a better identification of the attention/inattention states (Fox et al., 2016; Lutz et al., 2008). A meditation practice that involves focusing attention is Vipassana meditation characterized by focusing the attention on present-moment sensory awareness. In addition to improving the attention control, Vipassana meditation laid the basis for the development of mindfulness meditation techniques which have been used in clinical settings (Cahn and Polich, 2009; Goyal et al., 2014). Evidence suggests that such control of attention leads to changes in the functional neuroanatomical organization measured by modulations in functional connectivity (FC) (Hasenkamp and Barsalou, 2012; Kilpatrick et al., 2011; Rabipour and Raz, 2012).

FC is an estimation of the communication across distant brain regions, which results in the integration of information (Friston et al., 1993; van den Heuvel and Hulshoff Pol, 2010), a fundamental principle for cognitive processes. Various FC systems have been linked to attention tasks and FAM practices, primarily the frontoparietal network (FPN) (Bauer et al., 2019), the dorsal attention network (DAN) (Corbetta and Shulman, 2002; Raz, 2004; Vossel et al., 2014) and the default mode network (DMN) (Mooneyham et al., 2017). Focusing the attention to spontaneous sensations is also associated with changes in the functional connectivity of regions related to these three networks, according to previous studies (Bauer et al., 2014; Hasenkamp and Barsalou, 2012; Kilpatrick et al., 2011). However, most of these fMRI studies have shown results which implicitly considered FC as stationary, i.e. representing an average of the FC from the scanning session (Hutchison et al., 2013a; Preti et al., 2017), therefore, ignoring the dynamic character of the FC during attention processes and FAM practices (Fell et al., 2010; Hairston et al., 2017).

In this study we aimed at quantifying the dynamic variations of FC between DMN, DAN and FPN during three contiguous conditions; a resting state fMRI (rs-fMRI) session before a task, an spontaneous sensations attention task condition and during a resting state fMRI (rs-fMRI) session after the task, in experienced meditators. The inclusion of meditators provided the opportunity to explore the coordination of these brain networks during the interoceptive body focus in subjects with and advanced training in the control and sustaining of this process. (Lutz et al., 2008; Raffone and Srinivasan, 2010). The approach used in this study allowed us to determine and describe the characteristic FC patterns of the attention and interoceptive processes and its dynamics during the task and the resting state, as well as the determination of differences for this brain dynamics between the conditions, which has not been widely explored yet. We hypothesized that differences will be found in relation to the dynamic interaction of these three networks as an effect of the ongoing experimental condition.

## Materials and Methods

### Subjects

We included fifteen meditation practitioners in Vipassana meditation with an average number of hours of meditation practice at 1677 +/− 367, 6 females, with a mean age of 40 +/− 12 years old and. All participants were evaluated for exclusion criteria: fMRI contraindications, history of psychiatric or neurological disorder, or medical illness. Subjects answered digital versions of the Symptom Checklist 90 and Edinburgh Inventory to exclude psychiatric symptoms and to evaluate handedness, respectively. All participants signed an informed consent from the experiment approved by the Bioethics Committee of the Neurobiology Institute (Comité de Bioética del Instituto de Neurobiología, Universidad Nacional Autónoma de México).

### Experimental design

Functional images were acquired in one session for each of the 15 subjects. The session started with 10 minutes of resting state fMRI before the task (Pre-task rs), followed by 20 minutes of the attention task fMRI and finally 10 minutes of resting state fMRI after the task (Post-task rs) *(Fig. 1).* This implies three conditions for the subjects, nevertheless, one of the subjects did not conclude the post-task rs scan, therefore we obtained 14 subjects post-task rs data. The data for the pre-task rs and task condition was complete for the 15 subjects.

**Fig. 1.**
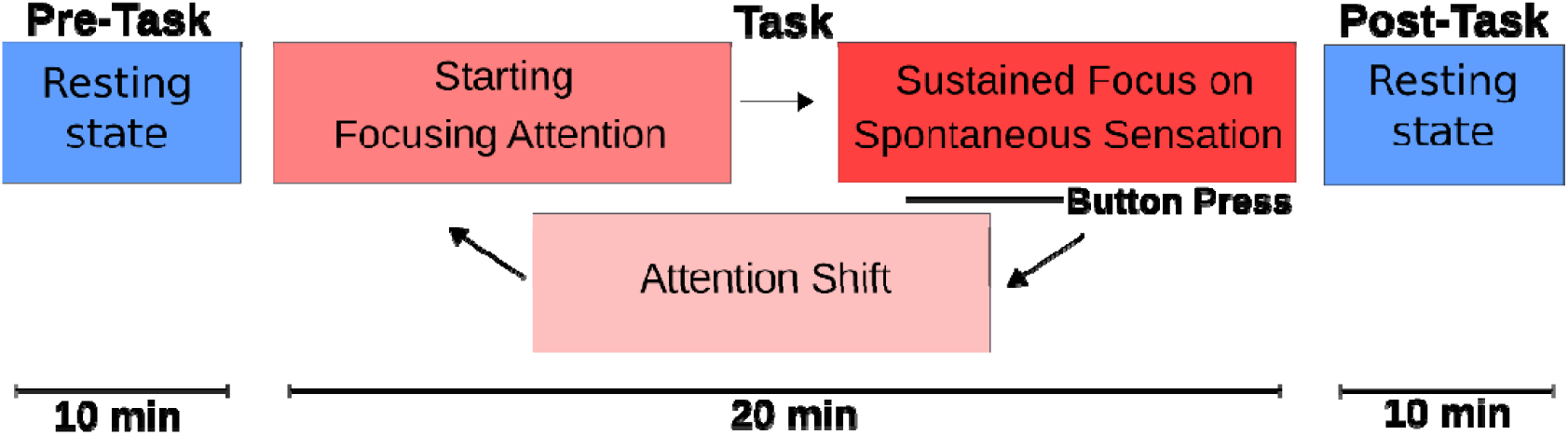
Experimental Design. The fMRI data were acquired in one session, starting with 10 min pre-task rs, followed by 20 min task scan and the 10 min post-task rs. The task consisted in focusing attention to spontaneous sensations starting in the nostrils. Once a clear spontaneous sensation was felt, subjects sustained the focus on it for 3-5 seconds. Then pressed a button to signal the shift of attention to the next target.

The attention task is a focus attention meditation technique, which consisted of focusing the attention to spontaneous sensations (e.g. numbness, pulsation, tingling, warming, cooling, itching, tickle, vibration, flutter, skin stretch, stiffness, etc.) in five specific anatomical regions: nostrils, right thumb, left thumb, right great toe, left great toe, always in the same order, cyclically and counterbalanced. Once participants started to feel a spontaneous sensation (i.e. pulsation) in the respective region, they were asked to sharpen focus more and more until they had a clear, distinct and uninterrupted sensation. When this level of felt sensation was reached, they were instructed to sustain it for about 3-5 seconds and then press a button (Nordic Neurolab MR compatible button system). This button press signaled both the end of clear and distinct focus of felt sensation and shift of attention to the next anatomical region *(Fig. 1)*. This was repeated at the participants own pace throughout the scan until the end. During the MRI session the subjects laid supine and remained relaxed with their eyes closed and avoided any motion. The time and the quantity of the responses were registered.

### MRI acquisition

Images were obtained using a 3.0T GE Discovery MR750 scanner (General Electric, Waukesha, WI) with a 32 channels head coil. We acquired three fMRI scans during one session per subject: A resting state scan before the task (pre-task rs), then, the task scan and finally a rest scan after the task (post-task rs). The attention task scan was obtained using a T2* EPI pulse sequence of 20 minutes, with TR/TE= 1500ms/27ms, 64×64 matrix, spatial resolution 4×4×4 mm3, 35 slices/volume, obtaining 804 volumes. The resting state fMRI scans consisted in an EPI pulse sequence of 10 minutes in duration each one, with TR/TE= 2000 ms/40ms, obtaining 300 volumes in the pre-task rs and 300 volumes in the post-task rs. During rest, subjects were asked to remain awake with their eyes closed. After the fMRI acquisition a high resolution 3D T1 SPGR structural sequence was acquired (voxel size of 1×1×1 mm3, 256×256 matrix, TR/TE= 8.156 ms/3.18 ms).

### ROI Selection

According to the literature, attention and focused attention meditation practices are associated with changes in functional connectivity in the DAN, DMN and FPN. Based on this evidence, we investigated the FC of these three networks. For this, we decided to use an individual parcellation approach, therefore, we created an individual mask in each subject containing the ROIs of the three networks, which would be used to make the FC analysis.

In order to create this individual masks, the pre-task rs data of each subject and the Functional Connectivity Toolbox (CONN) were used (Whitfield-Gabrieli and Nieto-Castanon, 2012). We performed a first level fMRI connectivity analysis in each pre-task rs data subject, after being preprocessed with a default preprocesssing of CONN; this included motion correction, slice timing correction, segmentation, coregistration to the MNI152 standard space, artefact detection, regression and spatial smoothing of 6 mm. For this analysis we used 4mm spheres, created with SPM in MATLAB, centered for the DMN, DAN and Executive control network (ECN) regions according to the coordinates of Raichle (Raichle, 2011). Then, we performed the voxel to voxel connectivity analysis obtaining a pFDR map for each network per subject. These maps were thresholded with a p<.05 value, then, to eliminate voxels out of our regions of interest that could survived to the threshold, we used FSLtools to multiply the thresholded binarized maps with the ROIs of the DMN, FPN/CEN and DAN of the network atlas implemented in CONN. This last step also allowed to obtain only the surviving voxels specific for each subject but within the boundaries of the CONN ROIs. The used CONN network atlas contains ROIs defined from CONNs ICA analysis in 497 subjects of the Human Connectome Project dataset (Whitfield-Gabrieli and Nieto-Castanon, 2012). After this analysis we observed that the number of ROIs for the DAN mask differ between the subjects, since for some subjects there were no significant voxels in one of the frontal eye fields. To homogenize the number of ROIs in the DAN masks, we resolved to contain only 2 regions, left and right intraparietal sulcus. The obtained three networks masks, per subject *(Fig. 2)*, were combined into one mask with 10 ROIs, subject specific. These consisted in the left and right intraparietal sulcus, the medial prefrontal cortex (MPFC), posterior cingulate cortex (PCC), left and right lateral parietal cortex (LPl, LPr), the left and right lateral prefrontal cortex (LPFCl and LPFCr), and the left and right posterior parietal cortex (pPCl, pPCr) *(Fig. 2)*.

**Fig. 2.**
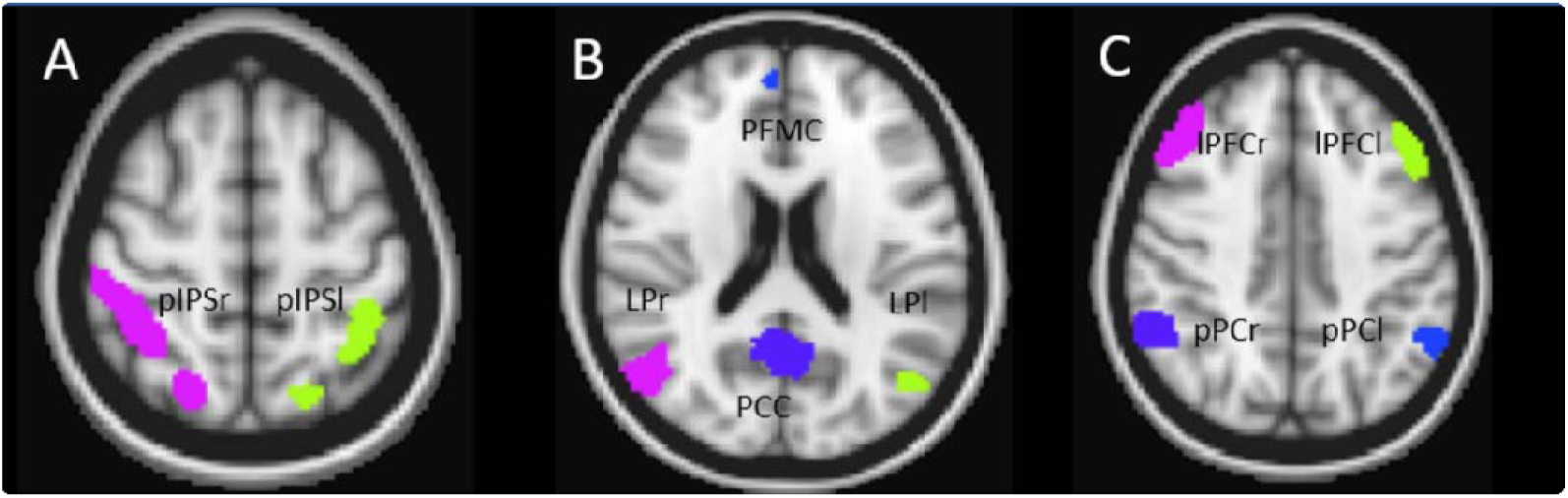
ROI masks. obtained from a subject including DAN (A), DMN (B) and FPN (C) regions. The 3 ROI masks were combined into a single mask, containing the 10 ROIs which was used for the DFC analysis of this subject.

### Preprocessing

After obtaining the individual ROI masks for every subject, the pre-task rs, task fMRI, and post-task rs fMRI raw data, were preprocessed using FSL (Smith et al., 2004) and for this the structural images were required. The steps for this preprocessing were: Extraction and discarding of skull and other non-brain tissue from the structural and functional image using BET of FSL (Smith, 2002) and reorientation. Slice timing correction, motion correction using MCFLIRT tool (Jenkinson et al., 2002), linear coregistration with FLIRT tool and nonlinear with FNIRT to the MNI152 standard space, segmentation of white matter and cerebrospinal fluid, regression of the signal from CSF, white matter and motion, artefact extraction with aCompCor (Behzadi et al., 2007) and band-pass filtering of 0.01-0.08 Hz. We did not perform the global signal regression since previous studies suggest that this may lead to false anticorrelations.

### Dynamic functional connectivity analysis

Sliding windows is a strategy applied to explore the time dynamic nature of the FC and in conjunction with a clustering approach as k-means, allows to identify patterns of FC that may reoccur in time across subjects, defined as dynamic functional connectivity states (DFCS) (Chang and Glover, 2010; Damaraju et al., 2014).

Applying a similar approach of previous studies (Damaraju et al., 2014; Mooneyham et al., 2017), we used sliding windows in order to determine the time varying FC (Sakoğlu et al., 2010) between the 10 ROIs of DAN, DMN and FPN in the preprocessed data (pre-task rs, task and post-task rs). Then, to determine the dynamic functional connectivity states (DFCS), we used the k-means algorithm for clustering (Allen et al., 2014; Chang and Glover, 2010) *(Fig.3).* For this DFC analysis we used FSL tools, R software (3.4 version) (R Foundation for Statistical Computing., 2016) and different R packages.

**Fig. 3.**
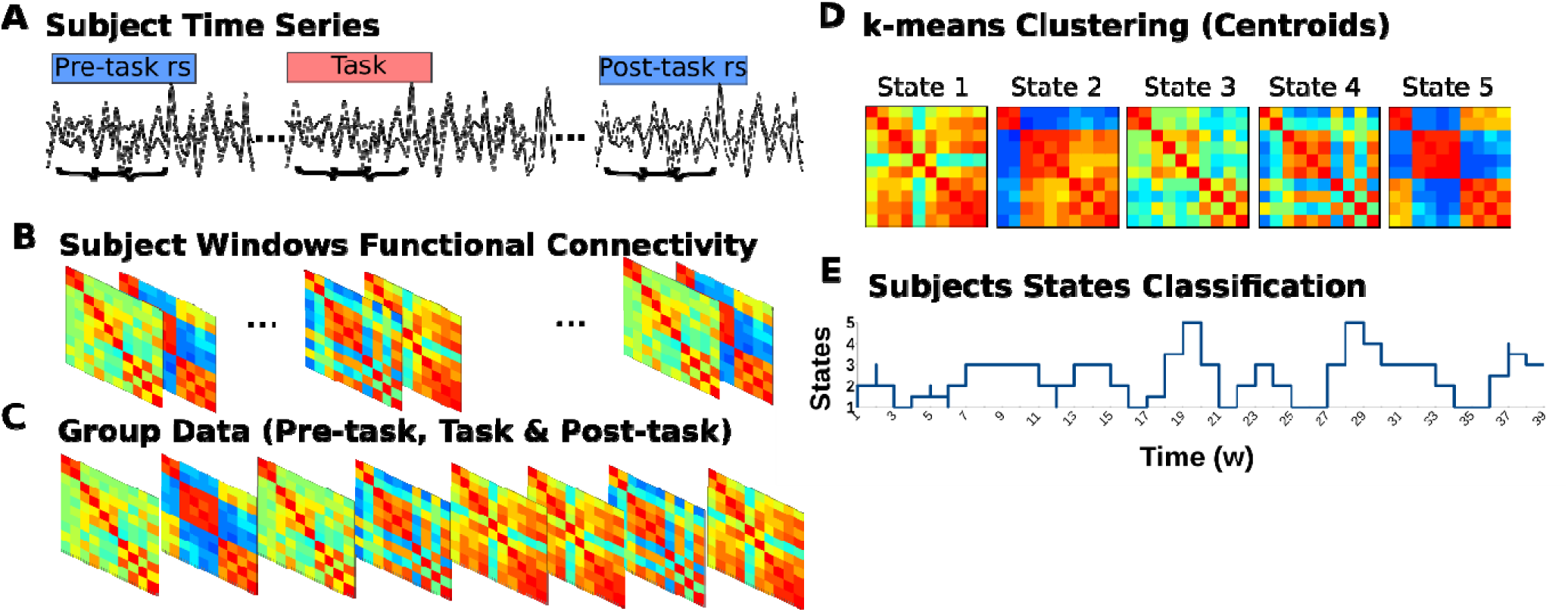
DFC Analysis. **(A**) The time courses of the three conditions in each subject where used for the sliding windows analysis. **(B)** In each 30s window the FC of the 10 ROIS was calculated. **(C)** The data from all subjects and conditions were joined into a group data set. **(D)** The cluster solution of 5 was applied and k-means clustering was used to obtain the centroids of each cluster, that represented the DFCS. **(E)** Each window (w), that represents 30s of time, was classified to one of the states, allowing to estimate the transitions and the time spent in the states for each subject and condition.

### Sliding windows and clustering approach

#### Sliding windows

For the sliding windows approach we established 30s windows width and 15s steps using fslsplit and fslmerge tools in the preprocessed data. The selected window size is according to previous studies describing a minimum windows size of 30s recommended to capture the FC fluctuations (Hutchison et al., 2013b; Tagliazucchi and Laufs, 2015). Therefore, in the task data we obtained 79 windows of 30s (20 TRs) width and 15s (10 TR) steps per subject, representing the 20 minutes of the subject task scan. This results in 1185 windows for the task group data. For the rest data, using 30s (15 TR) windows width and 15s (7TR) steps we obtained 39 windows for the 10 min of the pre-task rs and 39 windows for the 10 min of the post-task rs per subject. Therefore, in the pre-task group rs data we obtained 585 windows and in the post-task group data 546 windows of 14 subjects were obtained, since one subject did not conclude the post-task rs scan. The group windowed data, including the pre-task, task and post-task, consisted in 2,316 windows. In each window we calculated the functional connectivity between the ten regions of the three networks using the ROI mask estimated for each subject. For the FC estimation we computed the Pearson’s R for each pair of ROIs, obtaining a 10×10 matrix of connectivity for each window, then these values were transformed to z values.

#### Clustering

The 2,316 FC matrices containing z values were grouped into a data set that would be used to apply the k-Means clustering. To determine the ideal number of clusters for the grouped data set, we made an independent analysis. This consisted in determine the best number of clusters individually to the rest condition (pre & post-task rs) and to the task condition FC matrices using the NbClust package (Charrad et al., 2014) from R software. NbClust is a package that determines the optimal number of clusters in a data set using 30 indexes. According to the majority rule, the best number of clusters for the rest data was three clusters and for the task data was five clusters. Therefore, we decided to use the five cluster solution as the optimal number of clusters in the grouped data set (pre-task rs, task and post-task rs), allowing to fairly compare between conditions.

Applying the k-Means clustering in the grouped data we obtained the five centroids of the five clusters; these centroids represented the five dynamic functional connectivity states. The clustering analysis also allowed us to classify each FC matrix from the data set to one of these five states. Given that each matrix represents 30s of time, we estimated the proportion of time spent on each state for each condition per subject. For this, in each subject we determined the number of matrices classified to a DFC state during a condition, this number was divided by the total number of matrices of the condition; this resulted in a time proportion for each state and for condition per subject, which then allowed us to estimate differences in the proportion of time spent on a state between the conditions. In order to define if the experimental condition had an effect over the proportion of time spent in the FC states, we applied the mixed effects linear model using the lme function from the nlme package in R (Pinheiro, Bates, DebRoy, & Sarkar, 2017). This test was preferred since one of the subjects did not complete the 10 min of the post-task rs condition therefore we had a non-balanced sample. In the model fitting we used the proportion of time spent in each state by condition as the dependent variable and the condition and state interaction as the independent variable, with the subject by condition as the random effect. Then we tested the significance of the model effects with ANOVA. The post hoc comparisons were performed using the emmeans package from R software (Lenth et al., 2020) which obtains the estimated marginal means for the model and allows to compute the contrasts. The p values of the contrasts were then corrected for multiple group (states) comparisons with the Bonferroni test.

## Results

The dynamic functional connectivity analysis resulted in five DFC states, which are described here as states 1 to 5 and represent the pre-task rs, task and the post-task rs conditions *(Fig. 4B)*. Each state characterizes a distinct FC between the 10 ROIs of three networks, DAN, FPN and DMN, implicated in attention and focus attention meditation practices.

**Fig. 4.**
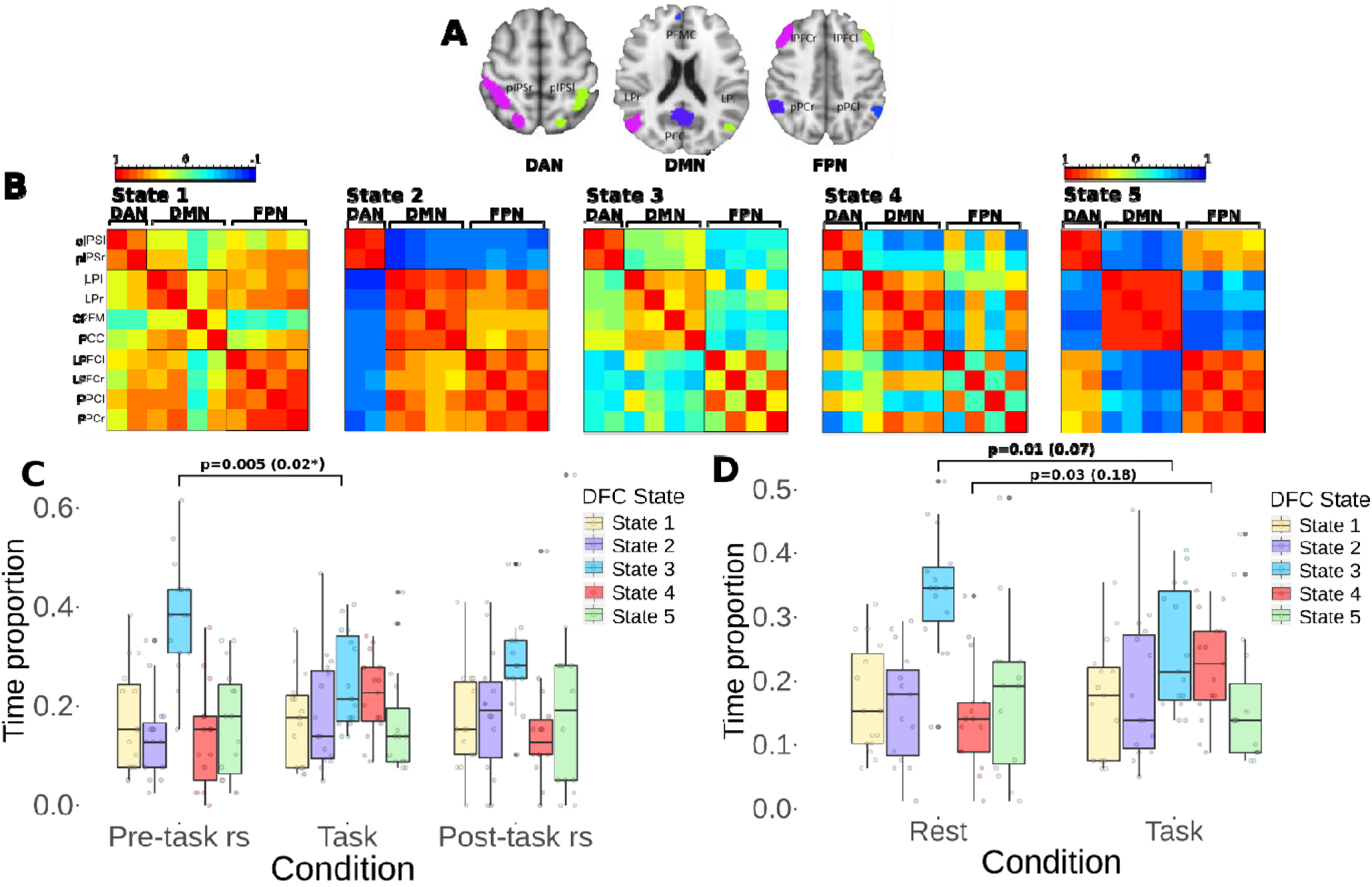
DFC States. **(A)**The individual masks where used for the analysis. **(B)** Five DFCS were obtained each one with a characteristic FC pattern and presented across the three conditions **(C)** Boxplots of the proportion of time spent in the each state shows differences between the three conditions. After being corrected, a significant decrease (*) in the time spent in the state 3 during the task performance in contrast with the pre-task rs was revealed. **(D)** An analysis contrasting between both rest conditions and task showed similar results, with a decrease in the time spent in the state three and an increase in the time spent in the state four, during the task, after the correction did not show significancy. DAN Dorsal attention network, DMN default mode network, FPN frontoparietal network.

The state one is characterized by a high connectivity between the regions of the three networks, except for the medial prefrontal cortex in the DMN, which shows low correlation values, near to 0, with the DAN and FPN regions. This state is also characterized by a preserved intranetwork connectivity. The state two shows a high connectivity between the DMN and FPN regions whereas, the regions from these two networks are anticorrelated with the DAN regions. Although the DAN is segregated from the two other networks, this shows a high intranetwork connectivity in this state. The state three demonstrates the lowest functional connectivity between the regions of the three networks in comparison with the other states. While this distinctive low internetwork connectivity points out a remarkable difference with the other states, it is also characterized by a preservation of the intranetwork functional connectivity. The state four is marked by the right FPN regions showing positive correlation values with the DMN regions, whereas, the left FPN regions show positive correlation with the DAN regions. This distinctive finding was associated to a decoupling between the left and right regions in the FPN, showing extremely low connectivity values between them. The state five shows a high connectivity between the DAN and the FPN regions, whereas, these two networks show anticorrelations with the DMN regions. Although this state shows a segregated DMN, this network exhibits the highest intranetwork connectivity in comparison with the other states.

The individual parcellations used for this study aimed to obtain the functional network regions specific for each subject, however, the use of a common anatomical atlas to study group FC is an approach widely used in fMRI (Salehi et al., 2018) Therefore, we decided to assess if the DFCS described in our study correlate with the states obtained using a common anatomical atlas, the CONN network atlas, for the group. We performed the same DFC analysis but using the ROIs of this atlas; these ROIs were the left and right intraparietal sulcus to study the DAN, the medial prefrontal cortex, posterior cingulate cortex, left and right lateral parietal cortex for the DMN and the left and right lateral prefrontal cortex, left and right posterior parietal cortex for the FPN. After the clustering approach using a five clusters solution, we obtained the five DFCS and visually compare these correlation matrices with the obtained using individual masks and it was observed that each of these five states had a matched state with one of the DFCS using the individual mask *(Fig. 5)*. A correlation Pearson’s test was performed between the z-values of the paired matching states, observing a significant high positive correlation between the five pairs of matching states. The results were for the State 1 r= 0.96, p=< 2.2^e-16^, 95% CI [0.93, 0.97], state 2 r= 0.98, p=< 2.2^e-16^, 95% CI [0.97, 0.99], state 3 r= 0.93, p=< 2.2^e-16,^, 95% CI [0.88, 0.96], state 4 r= 0.97, p=< 2.2^e-16^, 95% CI [0.95, 0.98], state 5 r= 0.99, p=< 2.2^e-16^, 95% CI [0.98, 0.99]]. This demonstrates a high degree of convergence between the functional connectivity states obtained using individual parcellations and using a group anatomical reference.

**Fig. 5.**
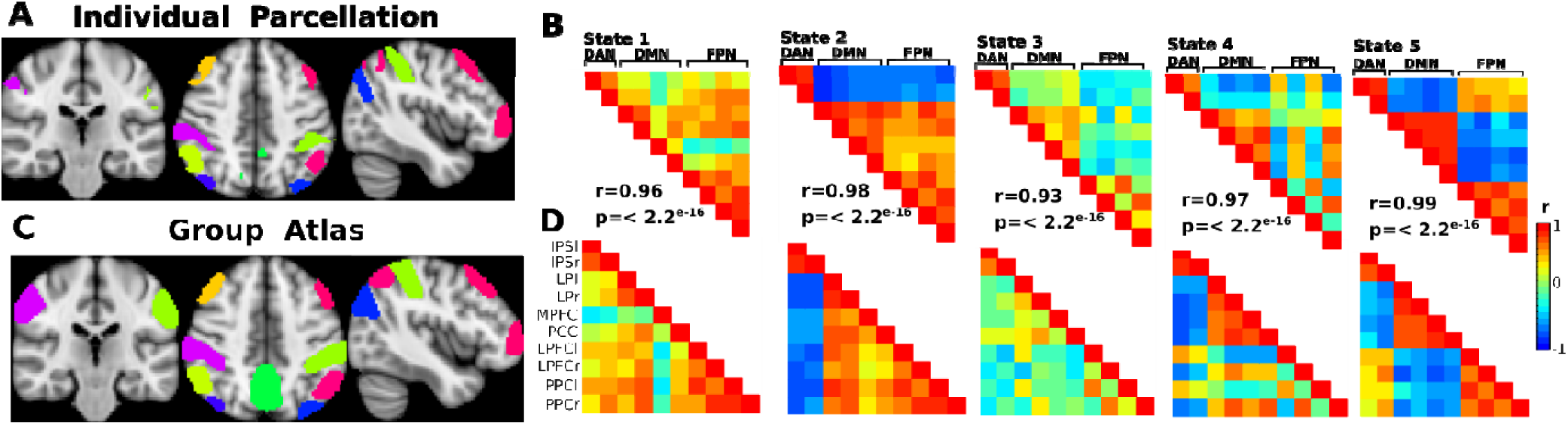
Comparison of the DFCS obtained with individual parcellations and with a common group atlas. **(A)**The ROI mask obtained from a subject is depicted. **(B)** These individual parcellations were used to obtain the five DFC states. **(C)**To asses the differences with a group common atlas, the 10 ROIS including DAN, DMN and FPN ROIS of the CONN network atlas was used. **(D)** Obtaining the five states of the common atlas. The correlation (r) and significancy between the visually matched states is shown. DAN Dorsal attention network, DMN default mode network, FPN frontoparietal network.

The application of the DFC analysis to the grouped data not only allowed us to characterize the DFCS that describe the brain dynamics of the three conditions, but also enabled us to investigate significant differences between the conditions. These differences were investigated in relation to the proportion of time spent in each of the five states for each condition.

The pre-task rs condition consisted in 585 FC matrices of the grouped data, which were classified each one to a DFCS. The analysis of the proportion of time spent in each state for this condition resulted in a mean proportion of 0.172 in the state one, 0.140 in the state 2, 0.381 in the state three, 0.138 in the state four and 0.167 in the state five. The task condition consisted in 1185 FC matrices that were classified to one of the five states. The calculation of the mean proportion of time resulted in a mean proportion of 0.172 spent in the state one, 0.188 in the state two, 0.251 in the state three, 0.221 in the state four and 0.165 in the state five during the task condition. For the post-task rs condition there were 546 FC matrices used to the analysis of the mean proportion of time spent on the states. This resulted in a mean proportion of 0.168 in the state one, 0.188 in the state two, 0.294 in the state three, 0.153 in the state four and 0.194 in the state five for this condition.

These results demonstrate differences in the proportion of time spent in each of the DFCS in relation to the condition, however, the differences were remarkable for the state three and state four. In the case of the state three there is an evident decrease in the time spent in this low functional connectivity state during the performance of the task in comparison to the pre-task and the post-task rs conditions. On the other hand, during the task performance there is a pronounced increase in the time spent on the state four characterized by a divided coupling of the FPN with the DAN and DMN, in comparison with the time spent in this same state during the pre-task rs and post-task rs conditions.

The statistical analysis using the linear mixed effect model and anova to the fitted model shows a significant effect of the condition and state interaction over the time proportion (F(8,191)=2.1473, p= 0.03), indicating that as predicted in our hypothesis, the proportion of time spent in the states is influenced by the condition. The post-hoc comparisons shows that there is a significant difference in the proportion of time spent in the state 3 between the pre-task rs and the task condition (p=0.005, 95% CI). After the correction for multiple group comparisons with Bonferroni test, we obtained a p=0.029 for this contrast. This indicates that there is a significant decrease in the time spent on the state three, the low connectivity state, during the performance of the task in comparison to the pre-task rs condition. There were no significant findings for the time spent in the other states in relation to the condition.

As is shown in Fig. 4C, differences are predominantly between the task and the rest conditions while both rest conditions share similar results. For this reason, we compared both rest conditions as a single rest condition with the task, to assess if the differences were more pronounced. The results show for the state 1 a mean proportion of time of 0.170 during rest and 0.172 during the task condition, for the state two a mean of 0.160 in rest and 0.188 during the task, the state 3 shows a mean of 0.341 in rest and 0.251 in task, the state four shows a 0.146 in rest and 0.221 during the task and the state five 0.182 during rest and 0.165 in task. The Fig. 4D illustrate these differences, which reflects once again a remarkable difference for the state three and state four between task and the rest condition. During the performance of the task there is a decrease in the time spent on the state three, while during this same condition an evident higher proportion of time spent in the state four is presented in comparison with the rest condition. The statistical analysis with the linear mixed effects model again demonstrates a significant effect for the state and condition interaction (F (4,126) =2.8486, p=0.026), indicating that the proportion of time spent in the states is affected by the experimental condition.

The post hoc comparisons between that rest-task contrast of the time for the state three resulted in p=0.014 and for the state four resulted in p=0.037 using a 95% CI. However, after the Bonferroni correction for multiple groups comparisons we obtained a p=0.07 and p= 0.18 for the state three and state four, respectively. There were no significant findings for the time in the other states in relation to these two conditions.

## Discussion

Previous studies of interoceptive attention and cognitive training techniques that improve its control, as FAM (Farb et al., 2013), have shown connectivity changes related to these processes, however the approach used in these studies assumed a constant FC in their results. Recent functional connectivity studies have shown the dynamic nature of the brain activity at rest, demonstrating FC patterns that evolve in time and even organize in a hierarchical way (Demertzi et al., 2019; Hutchison et al., 2013a; Vidaurre et al., 2017). Therefore, during the attention performance, the modulation of these dynamics may be subjected to specific characterization as brain activity is related to the cognitive demands (Gonzalez-Castillo and Bandettini, 2017).

The objective of this work was to determine the DFCS of the DAN, FPN and DMN regions during three different contiguous conditions, a pre-task resting state, an attention to spontaneous sensations task and a post-task resting state, as well as to effectively compare differences in the brain dynamics between these conditions.

Our main finding was that there are five different patterns of FC characterizing the conditions, each state with a specific connectivity between the three networks. A dynamic transition between the states was presented over the course of the three conditions, consistent with previous work stating that a varying brain FC configuration is essential for changes in behavior and cognitive demands (Tagliazucchi and Laufs, 2015). Moreover the proportion of time spent in each state was significatively related to the ongoing experimental condition confirming our hypothesis and in line with previous evidence that suggest that the time spent in these states is not random and that it might be associated with behaviour (Vidaurre et al., 2017). These differences were more evident between the task and the resting conditions, with the task showing a more complex organization in agreement with studies suggesting that task FC represents a combination of spontaneous activity and the task-related responses (Fox and Raichle, 2007; Fransson, 2006).

During the pre-task rs condition these five states were present, consistent with previous studies showing that brain activity during resting conditions is not abolished, instead, it was characterized by a varied set of FC patterns. However, the subjects spent a significant higher proportion of time in the state three in comparison with the task performance. This demonstrate that although a dynamic interaction between the networks was present, the tendency was to remain longer in a low internetwork connectivity brain pattern at rest.

This low connectivity state was the more prevalent across the three conditions, however, during the task there was a significant decrease in the time spent in this state. Considering this findings and the evidence that demonstrates that the time spent in these states is associated with the information processing (Vidaurre et al., 2017), we suggest that this low connectivity state is a basal and a transitioning state from which a state could transition to others. A similar pattern was found in an earlier DFC study which obtained three states including the executive control network (CEN), salience network (SN) and DMN regions, during a breathing attention task. One of the states of this study showed low correlations between the regions of the three networks and a preservation of the intranetwork arguing that this could represent a transitioning state (Mooneyham et al., 2017). In addition, the task condition was associated to a higher proportion of time spent in the state four, in comparison to both rest conditions, with a trend towards significance. This state showed the right FPN regions interacting with the DMN and the left FPN regions with the DAN and a decoupling of the left and right regions in the FPN. The flexible interaction of the FPN with DAN and DMN of this state is consistent with evidence showing that this sequential interaction is fundamental for the control and adaption in task demands (Cole et al., 2013; Harding et al., 2015). Although the FPN has been often viewed as a unitary system, the decoupling between its the left and right regions is consistent with recent works showing two susbsystems as an internal organization for this the network, one connected with the DMN and the other with DAN, which contributes to the ability to deliberately guide actions based on goals (Dixon et al., 2018). Moreover, the characteristics of this state agrees with previous meditation studies demonstrating that activity in FPN regions effects on the connectivity of the DMN (Bauer et al., 2019). This led us to suggest that this is a fundamental pattern of connectivity for the focused attention performance, carried out in this group of meditators. Although not significant, the task performance was associated to an increase in the average time spent in the state two, distinctive by a strong connectivity between DMN and FPN regions with segregation of the DAN. While the FPN and DMN are usually assumed to work in opposition (Crockett et al., 2017; Hsu et al., 2014), evidence suggest that their interaction may be involved in the regulation of introspective processes independent from sensory input and spontaneous thoughts (Christoff et al., 2009; Fox et al., 2015). The regulation of this introspective processes is a key characteristic of the focus attention meditation practices which would be consistent with the higher proportion found for this state during the sustained attention to spontaneous sensations.

The post-task rs condition was characterized by similar proportions of time spent in the states in relation to the pre-task rs. However, an exception was found for the state five, finding an increase in the average time spent in this state even though both rest conditions shared the same cognitive instructions.

This state showed a strong connectivity between DAN and FPN, networks that have been associated to focused attention states in meditation (Madhyastha et al., 2015; Mooneyham et al., 2017; Tang and Posner, 2009). In the case of the task against rest comparison, there was an increase in the time spent in this state for the task. Although not significant, this findings suggest a task-mediation effect extending to the post-task connectivity, in line with previous work showing that the rest functional connectivity succeeding a task, is affected by the prior cognitive state (Waites et al., 2005).

Over the three conditions, the state 1 characterized by a high connectivity between all the regions and a decoupling of the MPFC from the DAN and from the FPN was present in a similar mean proportion. MPFC supports self-related processing, emotional adaptive responses (Euston et al., 2013; Jang et al., 2011) and its connectivity is associated with conscious awareness (Luo et al., 2017). With regards to meditation, studies have found MPFC activity linked to mind wandering during focused attention to breath (Hasenkamp et al., 2012; Hasenkamp and Barsalou, 2012). Therefore, these results and the comparable distribution across the conditions lead us to suggest this could represent a mind wandering state, where subjects experience non directed thoughts.

In light of the findings and the previous evidence, we consider that the states found in our study represents the dynamic functional organization related to the cyclical attention states presented across the rest and the task performance (Tagliazucchi and Laufs, 2015). Beyond the description of the attention FC, our results could represent the dynamic integration of these networks in relation to interoception and body processing due to the task characteristics, with the permanence in each of the states as a fundamental way to efficiently fulfill the cognitive demands. The association of the interoception processing to a wide coordination between cognitive control networks suggest a top-down control from these networks to primary cortical regions linked to interoception, such as the insula and somatosensory cortex (Bauer et al., 2014; García-Cordero et al., 2017). This conclusion is supported by anatomic findings indicating inputs from parietal and occipital cortices to primary cortical regions which in turn integrates somatosensory information (Namkung et al., 2017).

Limitations of the study included a small sample size; however, this limitation was attenuated using an approach that allows to obtain a large amount of data points for the FC in each subject, that were included for the statistical analysis. The lack of a control group difficult the generalization of the results, this encouraged the design and the comparison to the resting condition. We realize the control group would enable the conclusions in a standard population, therefore, for future directions we propose to address these implications and additionally to include primary regions associated to interoception for characterize the brain coordination related to this interoceptive processes.

## Conclusions

In conclusion our study shows that both in rest and during the performance of an attention to interoception task, a time varying functional organization between the DMN, the DAN and the FPN regions is present. However, our results suggest that at rest, a restriction to remain in a low brain coordination is preferred. In contrast, a more complex brain integration with an interaction of functionally opposed networks is favored for the interoceptive process. Furthermore, it is interesting to observe the impact of the task performance over the subsequent resting FC and leads us to question if the time that this subsequent effect remains in FC could be related with the underlying mechanism of the learning process linked to meditation.

## Acknowledgments

AYM thank the “Programa de Doctorado en Ciencias Biomédicas, Universidad Nacional Autónoma de México (UNAM)” and the fellowship 721415 received from “Consejo Nacional de Ciencia y Tecnología (CONACYT)”. We are grateful to M.Sc. Leopoldo González-Santos and Dr. Erick Pasaye for technical support.

